# Mosquitoes do not like bitter: new perspectives for biorational repellents

**DOI:** 10.1101/2023.09.21.558838

**Authors:** Claudio R. Lazzari, Isabel Ortega-Insaurralde, Jérémy Esnault, Eloïse Costa, José E. Crespo, Romina B. Barrozo

## Abstract

Chemical repellents play a crucial role in personal protection, serving as essential elements in reducing the transmission of vector-borne diseases. A biorational perspective that extends beyond the olfactory system as the classical target may be a promising direction to move. The taste system provides reliable information regarding food quality, helping animals discriminate between nutritious and potentially harmful food sources, often associated with a bitter taste. Understanding how bitter compounds affect feeding in blood-sucking insects could unveil novel molecules with the potential to reduce biting and feeding. Here, we investigated the impact of two naturally occurring bitter compounds, caffeine and quinine, on the feeding decisions in female *Aedes aegypti* mosquitoes at two distinctive phases: (1) when the mosquito explores the biting substrate using external taste sensors and (2) when the mosquito takes a sip of food and tastes it using internal taste receptors. We assessed the aversiveness of bitter compounds through both an artificial feeding condition (artificial feeder test) and a real host (arm-in-cage test). Our findings revealed different sensitivities in the external and internal sensory pathways responsible for detecting bitter taste in *Ae. aegypti*. Internal detectors exhibited responsiveness to lower doses compared to the external sensors. Quinine exerted a more pronounced negative impact on biting and feeding activity than caffeine. The implications of our findings are discussed in the context of mosquito food recognition and the potential practical implications for personal protection.

## INTRODUCTION

Mosquitoes transmit disease to humans while fulfilling their reproductive requirements: anautogenous females must feed on vertebrate blood to complete their gonadotrophic cycles. Blood-feeding insects search for a host over long and short distances, mainly using their olfactory, thermal, and visual systems (Barrozo et al 2017; Lazzari, 2009; Montell and Zwiebel, 2016). Once landed on a host, the gustatory system or sense of taste allows animals to predict the quality of food, favouring the consumption of nutritious food and avoiding the consumption of toxic food. The taste sense plays an essential role during two different phases of blood-feeding behaviour: (1) when the insect reaches the host skin and has to decide whether to bite or not (i.e. the host skin recognition phase); and (2) when it takes the first sip of a blood meal and has to decide whether the food is adequate or not (i.e. the ingestion phase) (Barrozo, 2019). Thus, insects evaluate the skin and blood quality separately. The external taste sensilla of mosquitoes, located on the labella and tarsi, evaluate the presence of specific molecules on the biting substrate or host skin. Later on, the internal taste sensilla, located in the cibarial or in the pharyngeal region of the digestive tract, evaluate the quality of the ingested food (Barrozo, 2019).

Nutritious foods are signalled by sugars, amino acids, and low salt concentrations, whereas potentially harmful foods are signalled by bitter substances, acids and high salt concentrations (Shrestha and Lee 2023). Many toxins have a bitter taste to humans and this type of substances are biologically relevant in animal-plant relationships, as many plants produce them to protect themselves from herbivore attacks (Luo et al 2023). Although there is no distinct chemical identity for bitter compounds, their biological action is the same: they induce rejection or aversive behaviour in many animals, including blood-feeding insects (Glendinning, 2007; Barrozo, 2019). In *Aedes aegypti* mosquitoes, bitter compounds have been reported to be detected by the labellar sensilla (Sanford et al., 2013) and to modulate the response of sugar and water neurons in *Anopheles gambiae* (Kessler et al., 2013). Furthermore, sugar feeding is reduced when combined with bitter compounds (Ignell et al., 2010), and blood intake can also be negatively affected (Dennis et al., 2019; Kessler et al., 2014). Bitter compounds also prevent biting and feeding in the blood-feeding bug *Rhodnius prolixus*, by activating antennal and pharyngeal gustatory neurons, respectively (Pontes et al., 2014).

The sensory and behavioural aspects of biting and feeding in humans are key topics in medical entomology. Specifically, the taste system presents an attractive target for disease vector control due to its role in eliciting stereotyped and innate avoidance responses to certain tastants. However, understanding how these insects evaluate food quality and ultimately decide whether to bite or avoid a host based on taste remains a less explored area of study. To gain further insight into the effects of bitter compounds on mosquito biting and feeding, as well as to assess their potential value as gustatory repellents, we studied the effects of two naturally occurring bitter compounds, caffeine, and quinine, on the feeding decisions of female *Ae. aegypti* mosquitoes at two distinctive moments: the skin recognition and ingestion phases.

## MATERIALS AND METHODS

### Insects

*Ae. aegypti* of the Bora strain (insecticide-sensitive) were reared in the laboratory from eggs obtained from INFRAVEC (IRD -Montpellier, France) at 26 ± 1°C and 60 -70% RH. After their imaginal moult, the mosquitoes were kept with free access to sugar (imbibed cotton) but never fed with blood. Females between 15- and 20-days post-moult were used for the experiments.

### Artificial Feeder Tests

The feeding responses of individual mosquitoes were assessed in an artificial feeder (Figure 1). A female was placed in the mosquito container close to the membrane of the feeder, made of stretched ParaFilm M. The feeder or blood container contained 2 ml of fresh heparinized sheep blood (obtained from INRA, Nouzilly, France). Two series of tests were carried out with this set-up. In the first series, the bitter compounds were added to the feeder membrane while the feeder contained only blood (referred to as Contact tests). In the second series, the feeder membrane was kept clean while the bitter compounds were added to the blood container (referred to as Ingestion tests).

**Figure 1.**
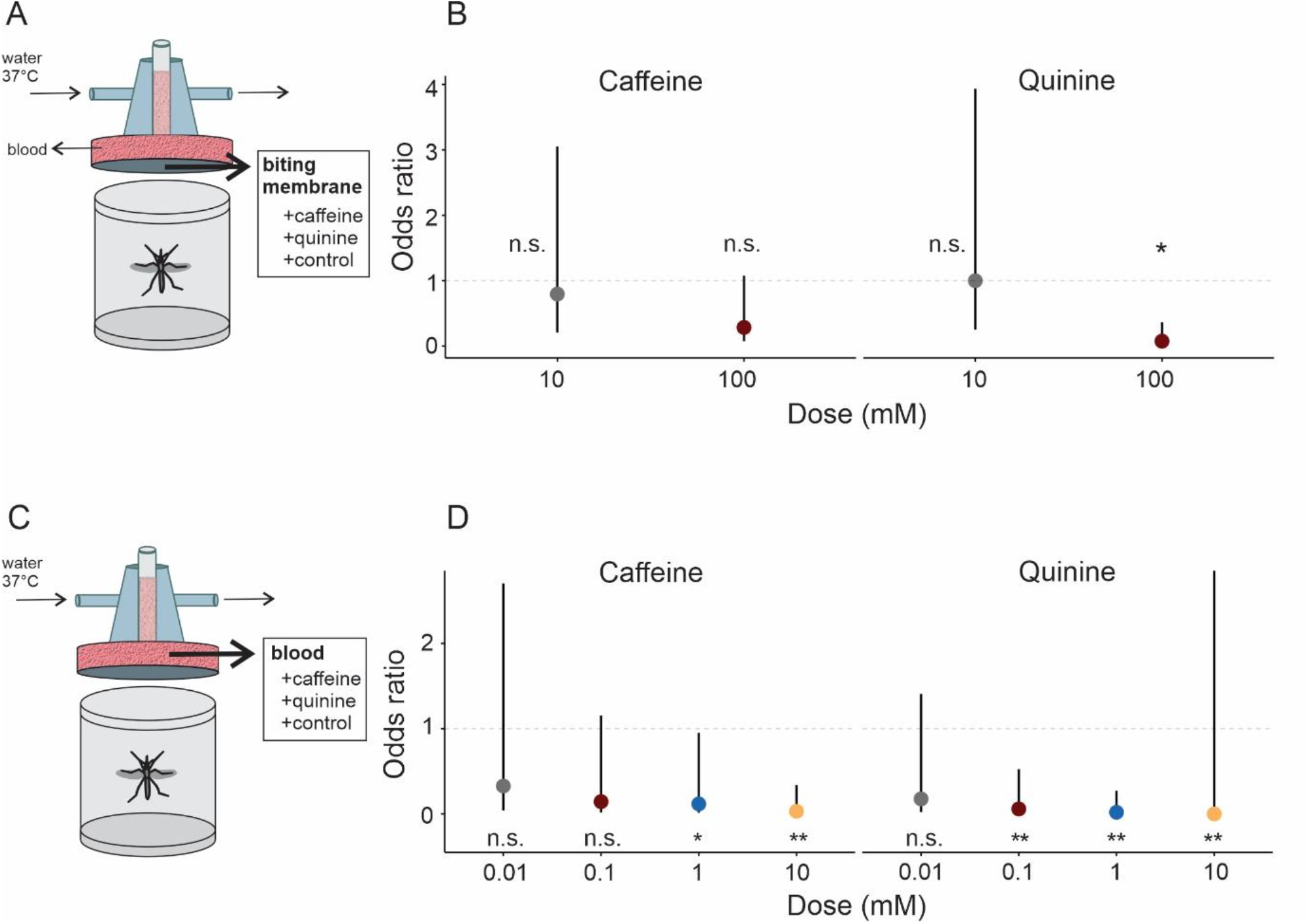
Suppression of feeding occurs upon detection of bitter compounds. (A) The experimental setup of the artificial feeder used for Contact tests is depicted. The biting membrane was coated with different doses of caffeine, quinine or distilled water (control), while blood was added to the feeder. (B) In Contact tests, we calculated the odds ratio, along with confidence intervals, to illustrate the likelihood of insects feeding on a membrane containing bitter compounds (caffeine or quinine) compared to insects coming into contact with the control membrane. The treatment with 100 mM caffeine led to a reduction in the probability of feeding among the treated insects, though this result was not statistically significant (Suppl. Table 1). Quinine, in contrast, produced significant reduction in the feeding probability at 100 mM compared respect to the control condition. (C) The experimental setup used for Ingestion tests is drawn. (D) In Ingestion tests, we determined the odds ratio with associated confidence intervals to illustrate the likelihood of insects ingesting bitter compounds (caffeine and quinine) compared to insects that ingested the control condition (blood). The treatment with 0.1 mM caffeine resulted in a reduction in the probability of feeding among the treated insects, although this finding was also not statistically significant (Suppl. Table 1). As the caffeine dose increased to 1 and 10 mM, the feeding response significantly decreased. Quinine induced a significant reduction in the feeding response starting from a dose of 0.1 mM. Statistical significance was assessed, and if the confidence interval for a particular comparison includes 1, it indicates that the odds ratio is not statistically significant, suggesting no discernible difference between the groups of insects. Significant differences are denoted by asterisks (* p < 0.05, ** p < 0.001), while ‘n.s.’ represents no significant differences. Each dose was tested in twenty replicates.

### Contact tests with bitter compounds

Twelve hours prior to the beginning of the experiments, feeder membrane surface (1.76 cm^2^) was coated with 100 μl of either 10 or 100 mM caffeine (CAS: 58-08-2, Biopack, Buenos Aires, AR) or quinine (CAS: 6119-47-7, Sigma-Aldrich, St Louis, US) dissolved in distilled water, resulting in doses of 0.00056 and 0.0056 mmol/cm^2^ on the membrane, respectively. A membrane coated with distilled water served as the control group. The behaviour of the mosquitoes was monitored for 10 min, after this period each female was assessed for feeding success by examining the visible swelling of the mosquito’s abdomen. Fed females were scored as 1, while unfed females received a score of 0. The score was measured as the visible swelling of the mosquito’s abdomen, in which case a success was recorded. Each individual was used only once and then discarded. Twenty individuals were tested per condition.

### Ingestion tests of bitter compounds

Caffeine or quinine was added to the blood in the feeder to achieve final concentrations of 0.01, 0.1, 1.0, or 10 mM. The blood-only feeder was used as a control. Mosquito behaviour was recorded for 10 min, after which each female was assigned a score of 1 if they had fed or 0 if they had not, based on the visible swelling of the mosquito’s abdomen, indicating a successful feeding. Twenty individuals were tested per condition.

### Arm-in-Cage Tests

To test for repellence of a real host, we adapted the standard arm-in-cage protocol recommended by the World Health Organisation (World Health Organisation, 2009). We made adjustments to the test to align with institutional security and ethical guidelines. Specifically, we ensured that the experimenter would not be bitten under any circumstances (even during control tests). Additionally, we adhered to the requirement that no substances causing allergic or toxic reactions in potentially vulnerable individuals should be applied to the skin. In addition, our protocol facilitated data collection and statistical analysis, resulting in balanced data between control and experimental tests. The experimenter wore heavy-duty latex gloves with 5 x 5 cm openings on the dorsal side of each hand (Figure 2a). A 6 x 6 cm piece of gauze was attached to the inside of each glove to cover the window. Another gauze piece, measuring 5.5 x 5.5 cm (30.25 cm^2^), was saturated with either the solvent (250 μl of distilled water) or the test solution (250 μl of a 100 mM solution of either caffeine or quinine), resulting in a 0.000826 mmol/cm^2^ on the gauze. The impregnated gauze was allowed to dry for 30 min before being inserted between the inner gauze and the window’s edges, where it remained. A 6 x 6 cm corrugated cardboard frame was positioned between the glove window and the hand, creating a barrier that prevented mosquitoes from reaching the experimenter’s skin. One hand of the experimenter wore the control glove, while the other wore the substance being tested. Each trial consisted in inserting one hand for 5 min, followed by a 30-min interval, and then introducing the second hand. In half of the tests, the control hand was introduced first, followed by the test hand. The order was reversed in the other half of trials. The number of landing mosquitoes on the gauze window and those biting attempts during the experimental period were counted.

**Figure 2:**
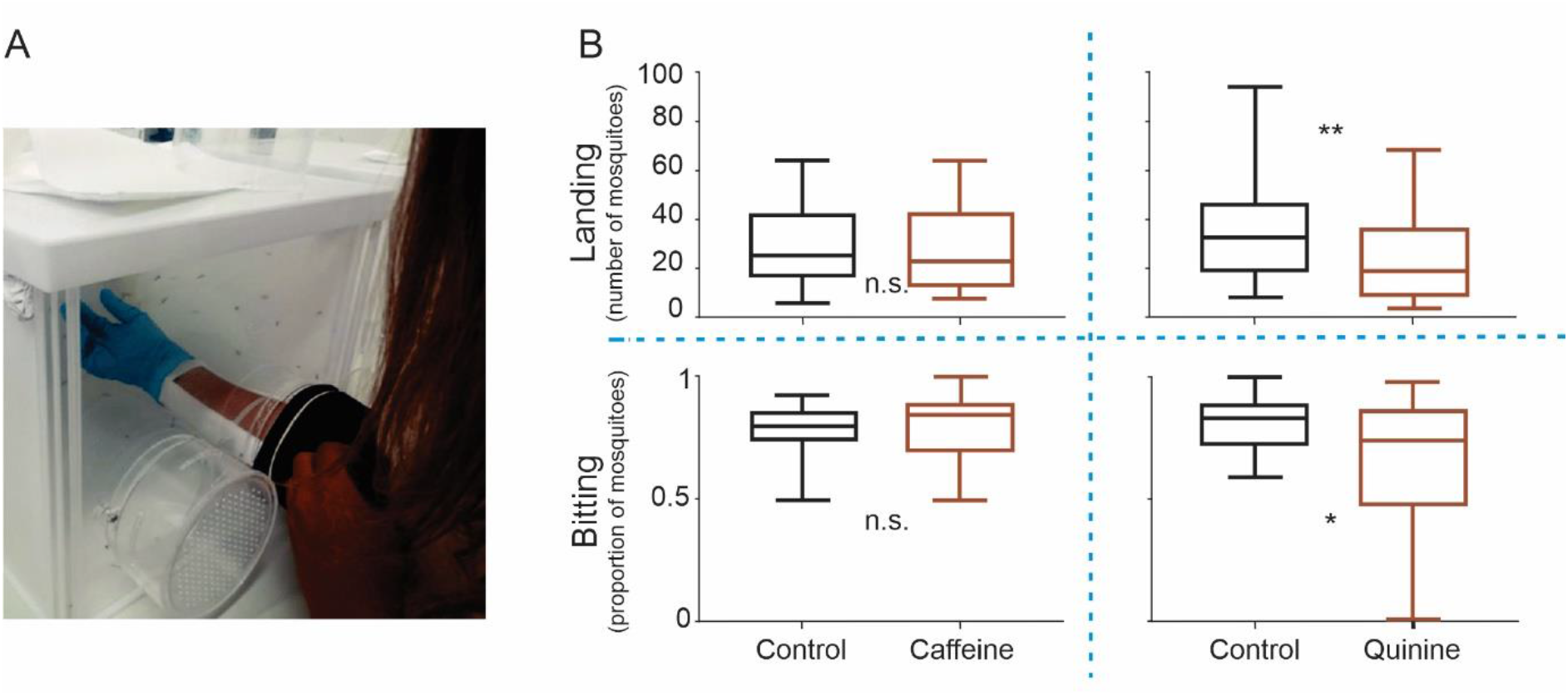
Reduced mosquito biting in response to bitter taste detection on a live host. (A) The arm-in-cage set up used in the experiment is shown. The experimenter’s arm is introduced into the mosquito cage while wearing a glove with an opening that has attached a gauze coated with 100 mM of caffeine, quinine or the solvent (control). (B) Top frames depict the number of mosquitoes that landed on both the treated and control hands of the experimenter. Bottom frames shown the proportion of landed mosquitoes that attempted to bite. Data are shown as median and 5-95 percentiles. Only quinine, but not caffeine, showed significant effect on landing or biting attempts with respect to the control hand. Significant differences are indicated by asterisks, (* p < 0.01; ** p < 0.001) while ‘n.s.’ denotes no significant differences. Eighteen replicates were carried out for each condition.

### Statistical Analysis

In the artificial feeder tests, we used a Binomial GLM with a logit link function to model the probability of feeding in individual *Ae. aegypti* mosquitoes during both Contact and Ingestion tests. The logit link function ensures fitted values between 0 and 1, and the Binomial distribution is typically used for probability data. The fixed factor was a combination of the bitter compound and different doses for both experiments. For the Contact tests, the categorical variables had five levels: control, 10 mM and 100 mM caffeine and quinine. For Ingestion tests, the categorical variables presented nine levels: control, 0.01 mM, 0.1 mM, 1 mM and 10 mM for caffeine and quinine.

We then used a Binomial GLM with a logit link function to compare the response to caffeine and quinine, assessing whether one bitter had a stronger effect than the other or if their effects were comparable. The fixed factors for Contact tests were bitter compounds (categorical with two levels, caffeine or quinine) and dose (categorical with two levels, 10 mM and 100 mM). The fixed factors for the Ingestion tests were the bitter compounds (categorical with two levels, caffeine or quinine) and dose (categorical with four levels, 0.01 mM, 0.1 mM, 1.0 mM and 1 0mM).

We performed all the analysis using the R v3.6.3 “Holding the Windsock’’ software. The glmmTMB and *nlme* packages were used to fit the models. We used the DHARMa package to test model assumptions. Plots were generated using the package *ggplot2*. Tukey contrasts were performed using the *emmeans* function of the *emmeans* package.

In the arm-in-cage experiment, we quantified the number of landings and the proportion of mosquitoes that attempted to bite on the control hand and the test hand. Neither of the two behavioural variables followed a Gaussian distribution across the groups. Therefore, we compared treatments using a Wilcoxon signed-rank test for paired samples (control hand vs. test hand).

## RESULTS

### Contact with a Bitter Membrane Suppresses Feeding

The aim of this series of experiments was to determine whether mosquitoes taste the host skin at the biting site and whether the presence of bitter compounds on the substrate could deter feeding (Figure 1a).

We found similar feeding probabilities at the low dose (10 mM) of caffeine and quinine as well in the control (Table 1). Both 10 mM caffeine (GLM, binomial, df = 95, p = 0.74) and 10 mM quinine (GLM, binomial, df = 95, p = 1) did not induce significant effects on feeding probability when compared to control insects that made contact with the untreated membrane (Figure 1b, Suppl. Table 1). These results suggest that mosquitoes do not detect the bitterness of these compounds upon initial contact at low doses. Furthermore, the mosquitoes exhibited consistent feeding behaviour during both the tests involving the control membrane and the tests with the low doses of bitter compounds.

**Table 1:**
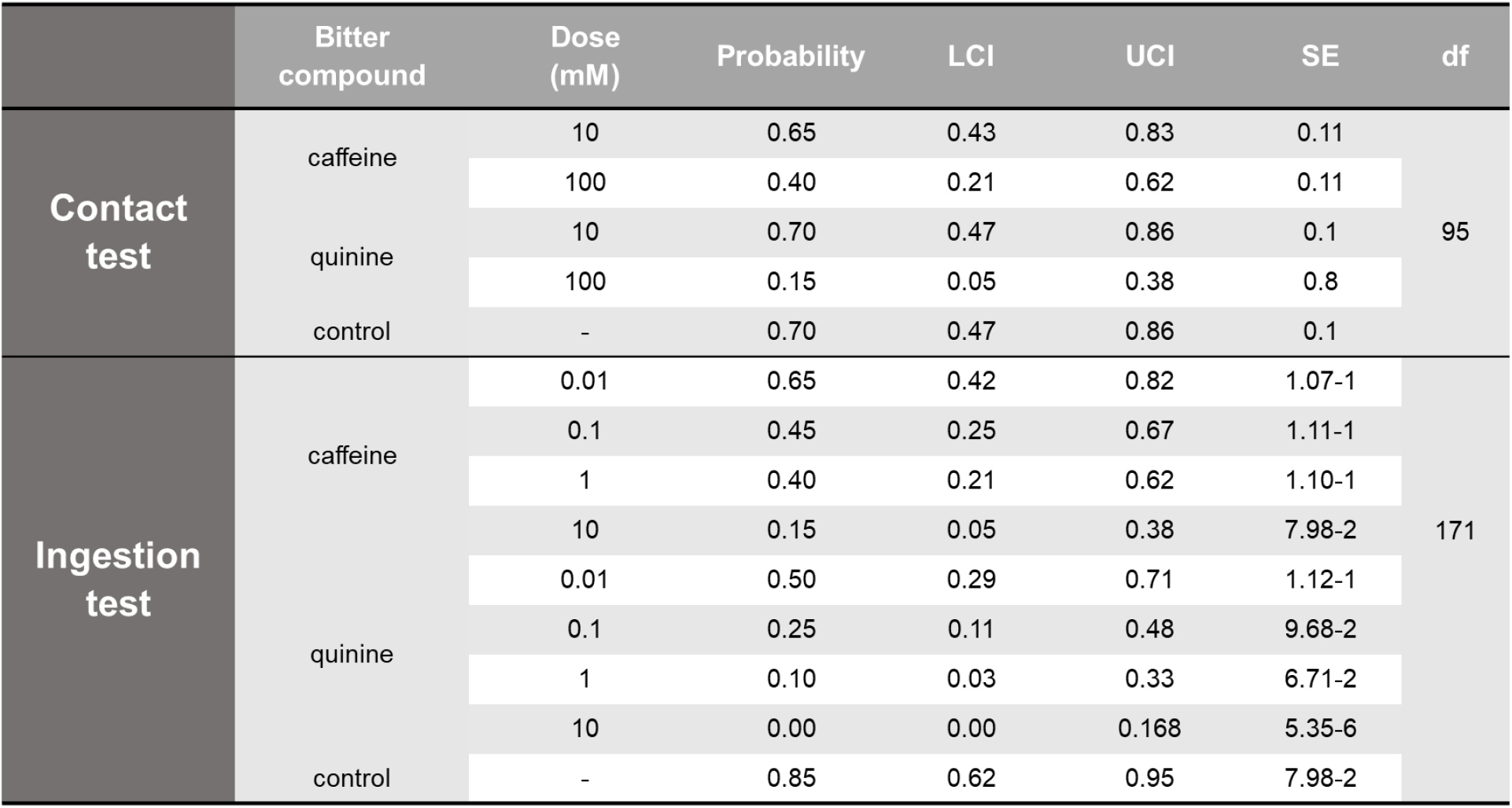
Summary of model results for estimating the feeding probability of individual mosquitoes. In Contact tests, the feeder membrane was coated with the bitter compound, while in Ingestion tests, the bitter compound was directly added to the blood. Control series served as the control group without any bitter compound. The “Probability” column represents punctual estimation, while “LCI” and “UCI” denote the lower and upper 95% confidence intervals, respectively. Abbreviations: SE = Standard Error; df = Degrees of Freedom.

However, at higher doses (i.e., 100 mM), we found a low but not yet statistically significant decrease in the probability of feeding for caffeine (GLM, binomial, df = 95, p = 0.06), and a marked and significant reduction in feeding for quinine (GLM, binomial, df = 95, p = 0.002) (Figure 1a, Suppl. Table 1). These results indicate that mosquitoes can detect high doses of bitter compounds, particularly quinine, leading to a change in feeding behaviour during the skin recognition phase. During the tests with the membrane coated with 100 mM quinine, we observed that the mosquitoes repeatedly made contact with the feeder membrane using their tarsi and labella but retracted their proboscis without piercing the membrane. Immediately afterwards, they moved to another area of the membrane and sampled again. This behaviour was consistently repeated, as if the insect were searching for a on the membrane without bitter taste.

Then, we compared the response of insects to caffeine and quinine to determine if one bitter had a stronger effect than the other or if their effects were similar. Although not statistically significant, we found that quinine had a slightly greater effect than caffeine but only at 100 mM (GLM, binomial, df = 76, p = 0.09) (Table 2). This result indicates that, starting from the 100 mM dose, the probability of biting is higher for caffeine than for quinine.

**Table 2:**
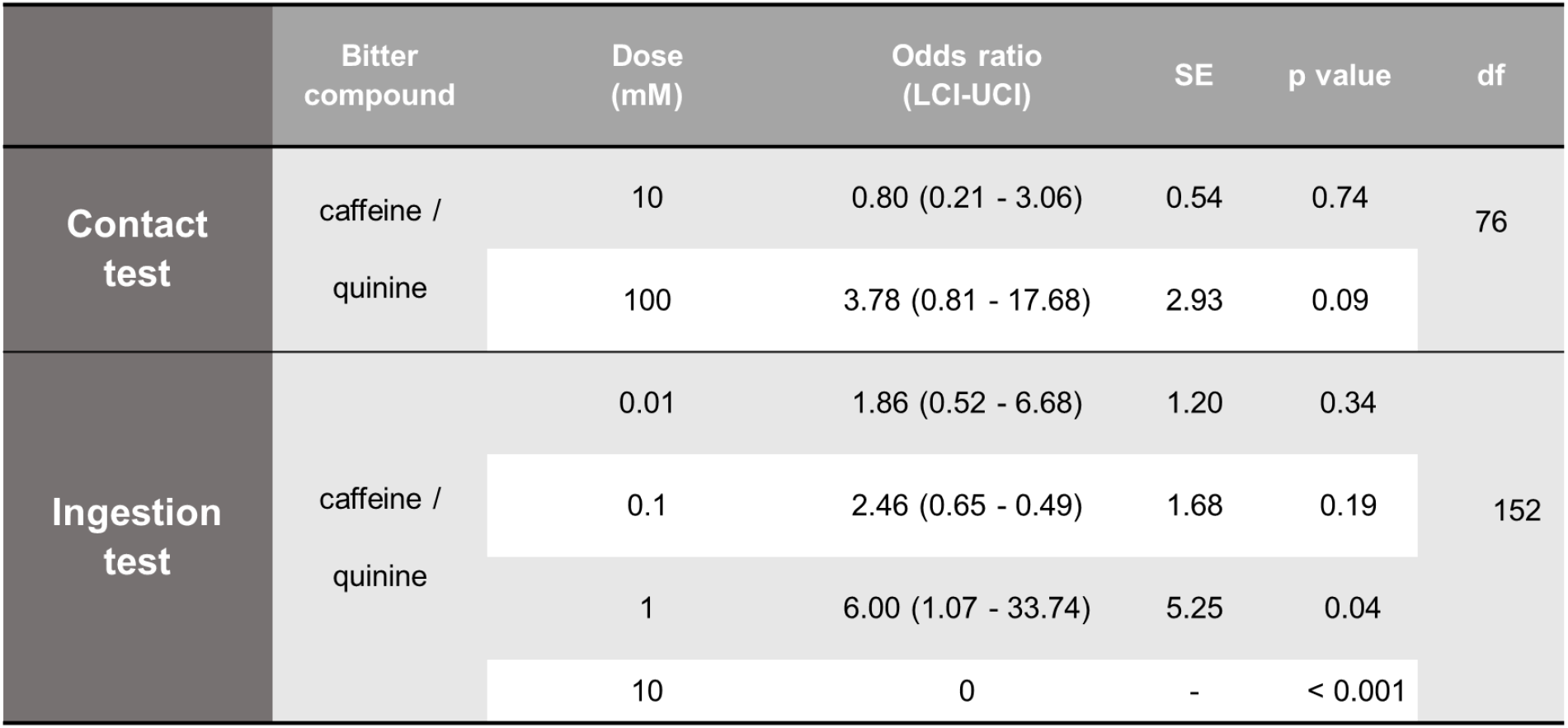
Odds ratios for bitter compounds at different doses in Contact and Ingestion tests. Odds ratios along with their corresponding 95% confidence intervals (LCI -UCI), comparing insects exposed to caffeine with those exposed to quinine in both the Contact and Ingestion tests are shown. Abbreviations: SE = Standard Error; df = Degrees of Freedom. Note that when the confidence intervals encompass the value 1, it indicates that there is no significant difference between the treatments.

### Perception of Bitterness in Food Deters Ingestion

Here, we asked whether mosquitoes could taste ingested blood, and whether the presence of bitter compounds in blood could inhibit feeding (Figure 1c).

At the lowest dose of caffeine or quinine (0.01 mM), we found similar feeding probabilities of both bitter compounds and the control (0.01 mM caffeine: GLM, binomial, df = 171, p = 0.58; 0.01 mM quinine: GLM, binomial, df = 171, p = 0.15) (Table 1, Figure 1d, Suppl. Table 1). However, starting from a dose of 0.1 mM, the feeding response decreased (Figure 1d, Suppl. Table 1). In the case of caffeine, at 0.1 mM, insects exhibited a tendency to reduce feeding compared to control insects, although not statistically significant (0.1 mM caffeine: GLM, binomial, df = 171, p = 0.08). As the dose of the bitter compound increased (1 and 10 mM), the feeding response further decreased (1 mM caffeine: GLM, binomial, df = 171, p = 0.04) (Table 1, Figure 1d, Suppl. Table 1). In contrast, quinine induced a significant reduction in the feeding response starting from a dose of 0.1 mM (0.1 mM quinine: GLM, binomial, df = 171, p = 0.005) (Figure 1d, Suppl. Table 1).

Similarly for Contact tests, we compared the response of insects to caffeine and quinine, aiming to ascertain whether one bitter substance exhibited a more pronounced effect than the other or if their effects were comparable. We found that quinine exerts a stronger effect than caffeine at high doses (1 and 10 mM) (1 mM: GLM, binomial, df = 152, p = 0.04; 10 mM: GLM, binomial, df = 152, p < 0.001) (Table 2). This result suggests that the impact of quinine is significantly greater than that of caffeine. Specifically, starting from a dose of 1 mM, the probability of feeding is higher for the caffeine than the quinine.

### Detection of Bitter Taste on a Real Host Reduces Biting

To assess the effect of caffeine and quinine in the context of a live host, we conducted arm-in-cage tests (Figure 2a). Figure 2b illustrates the results of these tests. Caffeine showed no significant effect on either landing or biting attempts when compared to the respective control condition (n.s.). In contrast, quinine produced a significant decrease in the occurrence of both landing (p < 0.001) and biting attempts (p < 0.01) when compared to the control situation.

## DISCUSSION

In this work, we investigated the effects of caffeine and quinine on modulating decision-making in the mosquito *Ae. aegypti* during host-skin recognition, and the ingestion phases. Interestingly, this effect was different between both bitters in terms of the behavioural impact observed. If we consider that the feeding process involves landing, skin tasting, biting and ingestion, it becomes evident that neither the effect nor the efficacy of these two compounds were consistent at the lowest doses used in our experiments.

The caffeine-treated membrane of the artificial feeder reduced mosquito feeding. While caffeine did not inhibit landing on a host’s hand (which is not surprising for a substance intended to be detected upon contact), it failed to prevent landing and biting when presented on a real host. However, it is worth to note that amount of mass of caffeine per cm^2^ applied on the real host was much lower in comparison to the artificial feeder tests. It is likely that mosquitoes require higher doses of caffeine to be repelled. However, when caffeine was directly incorporated into the blood, feeding was significantly reduced at relatively low doses. Quinine, on the other hand, was found to inhibit mosquito feeding behaviour at all tested conditions. Blood-feeding on the artificial feeder was significantly reduced when quinine was applied to the membrane or mixed with the blood. It was also effective in reducing landings and attempts to bite the host’s hand in the arm-in-cage tests. The latter effect is expected for a genuine contact repellent, but the former also suggests that quinine can be perceived before any physical contact with a treated substrate, possibly in a volatile phase. This finding suggests that either contact chemoreceptors detect it in the air, probably in close proximity where the concentration is relatively higher, or that olfactory receptors are also involved in the detection of quinine. In any case, our results confirm the concept that bitter compounds hold promising potential as contact repellents, which is worth exploring further.

Caffeine and quinine are likely to be first detected by the tarsi, particularly the fore- and middle-tarsi, during the host skin recognition phase (Dennis et al., 2019; Baik and Carlsson, 2020). Little is known about the sensory neurons within the tarsal trichoid sensilla of mosquitoes. The tarsal gustatory sensilla of *Ae. aegypti* are sex dimorphic, and contain up to five neurons, which have been classified into three categories (C1, C2, C3) (McIver and Siemicki, 1978). The electrophysiological responses of these sensilla to salts, sugars and amino acids have been mainly related to the oviposition and nectar-feeding behaviour (McIver and Siemicki, 1978). It is plausible that the previously reported tarsal neurons that respond to DEET on contact are the same that respond to high doses of quinine, caffeine and other bitter compounds (Dennis et al., 2019). After the mosquito lands on the substrate, the labellar lobes of the proboscis also contact the host skin. The medium-length hairs of the *Ae. aegypti* labella have been characterised previously (Sanford et al., 2013). One out of three neurons responded with small-amplitude spikes to both quinine and DEET. In contrast, caffeine, tested at 1 mM, failed to elicit an electrophysiological response that is consistent with our behavioural results (Sanford et al., 2013). In addition, a putative bitter receptor called AaegGr14, orthologous to the *An. gambiae* AgGr2, has been found in the labella transcriptome of *Ae. aegypti* (Sparks et al., 2013). Whether the detection in the labella is also responsible for the aversive behaviour towards the bitters on the feeding membrane is debatable, as Dennis et al. (2019) found that DEET detection was solely performed by the tarsal sensilla. Although not yet clearly demonstrated, a possible role for both mosquito tarsal and labellar sensory neurons would be the detection of bitter compounds, including those that may be present on the host skin, as well as defensive plant metabolites. In this way, a stage of contact repellency would operate in mosquitoes prior to the actual biting of the host skin, as has been demonstrated in kissing bugs (Pontes et al., 2014; 2022).

Although, we demonstrated that both caffeine and quinine present in the blood were aversive and dose-dependent during the ingestion phase, quinine elicited a higher aversiveness than caffeine at lower doses. The antifeedant effect of caffeine and quinine in the diet has already been demonstrated in *Ae. aegypti* and *An. gambiae* (Dennis et al., 2019, Ignell et al., 2010; Kessler et al., 2014). Interestingly, our results indicate that the presence of quinine in the blood led to a significantly higher level of aversion in *Ae. aegypti* compared to *An. gambiae* at the same concentration (Kessler et al., 2014). It is likely that receptors detecting bitter compounds in the blood are located in either the labrum or cibarium of mosquitoes (Lee, 1974; Lee and Craig, 1984). Both organs contain gustatory sensilla that could possibly be activated by the incoming blood. Neurons in the labrum are the first to detect blood (Jové et al., 2020). Four subsets of labral neurons detect specific components of the blood, including the nucleotide ATP, the salts NaCl and NaHCO3, and glucose (Jové et al., 2020). Cibarial neurons are also candidates for active detection of blood components, but their characterisation remains unknown. Bitter detection during feeding in the blood-feeding bug *R. prolixus* has been postulated to occur in the sensilla of the pharyngeal organ, which would function analogously to the cibarial organ of mosquitoes (Pontes et al., 2014).

Another interesting finding in this study is that bitter perception through two sensory pathways, i.e. external and internal gustatory sensilla, showed different acceptance thresholds. Taste perception of caffeine and quinine through the external taste receptors appears to be less sensitive than through internal taste sensors. In a previous study conducted in *Ae. aegypti* females, it was demonstrated that low doses of bitter compounds such as denatonium, lobeline, and quinine did not elicit feeding aversion when detected on the host skin (Dennis et al., 2019). However, these bitter compounds induced varying levels of significant aversiveness when added to the ingested food compared to the control. This prior work complements the results observed here, both in the context of external sensing and internal taste detection. It appears that the two instances of taste operate with different sensitivity filters, with the external one being less sensitive than the internal one.

Bitter compounds are known to cause negative post-ingestive effects in many different animals (Glendinning, 2007). The metabolic effects of ingestion of these toxic compounds can impair learning, change the behaviour or even cause death (Muñoz et al., 2020; Saunders et al. 1992). Some bitter compounds are both unpalatable and toxic to insects, while others are only unpalatable. Internal filters are the last preventative barrier to avoid deleterious physiological consequences. Therefore, taste-related barriers could protect animals from drastic consequences. Many of the substances currently used for personal protection, as DEET and other repellents, primarily target the olfactory system in the vapour phase, aligning with the classical definition of repellents (Barton-Browne, 1977). The crucial role played by the taste system in feeding decisions of insects has been known for a long time, making it an intriguing target for controlling the transmission of vector-borne infectious diseases. Despite progress in understanding the molecular targets of repellents, the search for new molecules is hindered by a lack of precise knowledge regarding their mechanism of action. We still do not fully understand why repellents effectively deter insects. Several hypotheses have been proposed, but none seems to account for all the experimental evidence, not even DEET, which has served as the gold standard for repellents for over sixty years. DEET’s mode of action is sometimes described as “enigmatic” or “mysterious” (Leal, 2014; DeGennaro, 2015). Nonetheless, chemical repellents for personal protection remain essential in mitigating the transmission of vector-borne diseases. A deeper understanding of these molecules and their impact on vector behaviour could open new perspectives in the search for a novel generation of repellents.

## ACKNOWLEDGEMENTS

Authors express their deepest gratitude to the institutions that made possible this study: CNRS, University of Tours, University of Buenos Aires, and CONICET,

## AUTHOR CONTRIBUTIONS

RBB and CL: Conceptualization, Resources, Supervision, Funding acquisition, Project administration, Data curation, Formal analysis, Investigation, Methodology, Validation, Visualization, Writing – original draft, Writing – review & editing. JE, EC: Investigation, Formal Analysis. IOI, and JEC: Data curation, Formal analysis, Software, Validation, Visualization, Writing – original draft.

## ETHICS STATEMENT

All the experiments were conducted respecting the hygiene and security policies of the involved institutions. Experimenters were preserved from eventual expositions to mosquitoes and chemicals employed.

## Figures and Tables

**Supplementary Table 1:**
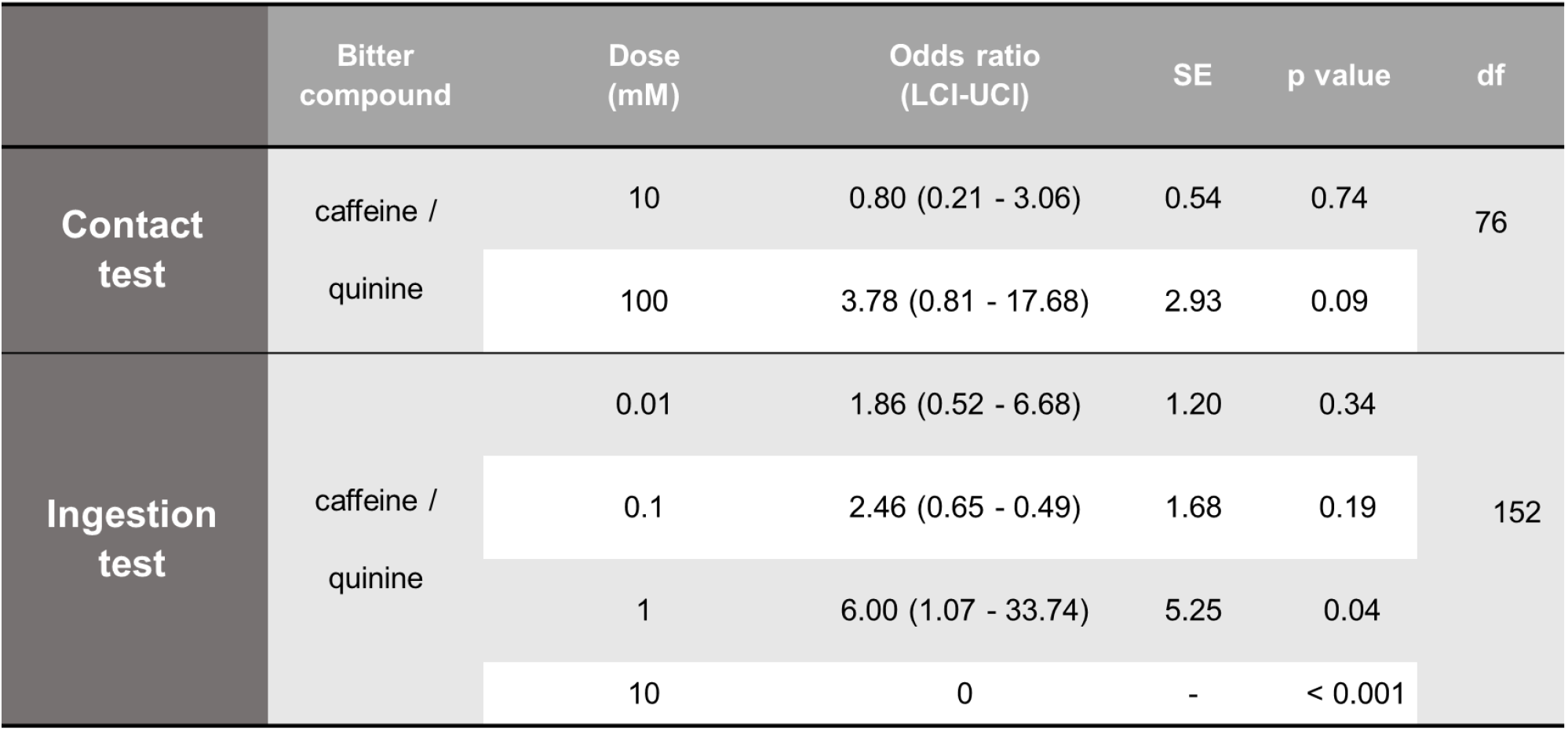
Odds ratios for bitter compounds at different doses in Contact and Ingestion tests. Odds ratios along with their corresponding 95% confidence intervals (LCI-UCI), comparing insects exposed to bitter compounds (caffeine or quinine) with control insects in both Contact and Ingestion tests are shown. Abbreviations: SE = Standard Error; df = Degrees of Freedom. Note that when the confidence intervals encompass the value 1, it indicates that there is no significant difference between the treatments.

